# Blocking chondrocyte hypertrophy in conditional *Evc* knockout mice does not modify osteoarthritis progression

**DOI:** 10.1101/2021.10.29.466392

**Authors:** Ana Lamuedra, Paula Gratal, Lucía Calatrava, Víctor Luis Ruiz-Perez, Adrián Palencia-Campos, Sergio Portal-Núñez, Aránzazu Mediero, Gabriel Herrero-Beaumont, Raquel Largo

## Abstract

**Background:** Chondrocytes in osteoarthritic (OA) cartilage acquire a hypertrophic-like phenotype, where Hedgehog (Hh) signaling is pivotal. Hh overexpression causes OA-like cartilage lesions, whereas its downregulation prevents articular destruction in mouse models. Mutations in *EVC* and *EVC2* genes disrupt Hh signaling, and are responsible for the Ellis-van Creveld syndrome skeletal dysplasia. Since Ellis-van Creveld syndrome protein (Evc) deletion is expected to hamper Hh target gene expression we hypothesized that it would also prevent OA progression avoiding chondrocyte hypertrophy. Our aim was to study Evc as a new therapeutic target in OA, and whether Evc deletion restrains chondrocyte hypertrophy and prevents joint damage in an Evc tamoxifen induced knockout (*Evc*^*cKO*^) model of OA.

**Methods:** OA was induced by surgical knee destabilization in wild-type (WT) and *Evc*^*cKO*^ adult mice, and healthy WT mice were used as controls (n=10 knees/group). Hypertrophic markers and Hh genes were measured by qRT-PCR, and metalloproteinases (MMP) levels assessed by western blot. Human OA chondrocytes and cartilage samples were obtained from patients undergoing knee joint replacement surgery. Cyclopamine (CPA) was used for Hh pharmacological inhibition and IL-1β as an inflammatory insult.

**Results:** Tamoxifen induced inactivation of *Evc* inhibited Hh overexpression and partially prevented chondrocyte hypertrophy during OA, although it did not ameliorate cartilage damage in DMM-*Evc*^*cKO*^ mice. Hh pathway inhibition did not modify the expression of proinflammatory mediators induced by IL-1 beta in human OA chondrocytes in culture. Hypertrophic – IHH – and inflammatory – COX-2 – markers co-localized in OA cartilage samples.

**Conclusions:** Tamoxifen induced inactivation of *Evc* partially prevented chondrocyte hypertrophy in DMM-*Evc*^*cKO*^ mice, but it did not ameliorate cartilage damage. Our results suggest that chondrocyte hypertrophy per se is not a pathogenic event in the progression of OA.

## INTRODUCTION

Osteoarthritis (OA) is a chronic joint disease mainly affecting articular cartilage, which undergoes erosion, characterized by extracellular matrix (ECM) degradation and cell alterations(1–5). Chronic biomechanical stress is the main factor triggering cartilage degradation(5). The resulting damage-associated molecular patterns (DAMPs) activate Toll-like receptors (TLR) and innate immunity in OA chondrocytes, evoking a local chronic inflammatory response with an increase in proinflammatory cytokines such as interleukin (IL)-1 or tumor necrosis factor (TNF). In turn, these cytokines induce the release of active metalloproteinases (MMP), aggrecanases and different proinflammatory mediators which activate the catabolic program characteristic of this disease(5). Together with inflammation, regenerative mechanisms are triggered by OA chondrocytes presumably as an attempt to repair the damagedtissue, such as the reactivation of signalling pathways operating during endochondral ossification of the growth plate, as Indian Hedgehog (IHH), WNT or NOTCH signaling(1,6,7). That is the reason why OA chondrocytes with this gene expression pattern are known as hypertrophic-like chondrocytes.

During limb development hypertrophic chondrocytes show a gradual increase in its cell size and varying gene expression, including Runt-related transcription factor 2 (RUNX2) and IHH, and progressively type X collagen (COL-10), MMP-13, receptor activator of nuclear factor kappa-B ligand (RANKL), and osteopontin (SPP1)(1,8). This developmental process, deeply coordinated by the IHH–parathyroid hormone-related protein (PTHrP) axis, drives an active ECM remodeling, until chondrocytes reach an apoptotic fate and leave a mineralized matrix for bone formation(1,8). Therefore, IHH has an essential role during embryonic and postnatal skeletal development and bone growth.

The Hedgehog (Hh) family of signaling molecules mediates the development of numerous organs during embryogenesis. However, In general, this signaling is physiologically repressed in the adult stage being only reactivated during tissue repair processes after injury, such as in lung epithelium, muscle, cartilage and bone(9).

Three Hh genes have been described in vertebrates, Sonic (SHH), Desert (DHH) and IHH, whose canonical signaling anchors to the primary cilium, a cellular structure highly specialized in the reception and transduction of mechanical and biochemical stimuli into the cell(10). Upon the arrival of a Hh ligand and its binding to the PTCH1 receptor, Smoothened (SMO) is released from PTCH1-mediated repression, and it translocates to the base of the cilium, where interacts with the EVC-EVC2 complex(11,12). This interaction promotes the activator function of the gliome associated oncogene (GLI)transcription factors and thus the expression of Hh target genes, including Hh signaling components, such as PTCH1 and GLI1, and chondrocyte hypertrophy related genes(2,11–13).

Congruently with the hypertrophic-like phenotype, both human and animal OA cartilage exhibits increased levels of IHH and overexpression of components of the Hh pathway(14,15). Furthermore, chondrocytes in OA cartilage have also been described as morphologically hypertrophic cells with an increased size that are present in all the articular cartilage zones(16). These cell alterations also correlate with the expression of IHH in the cartilage and with the grade of tissue destruction(13,15,17). Experimental models have revealed that genetically modified mice with higher activation of Hh signaling (*Ptch1*^*+/−*^, *Col2a1-Gli2*– transgenic and *COL2-rtTA-Cre;Gt(ROSA)26Sor*^*tm1(Smo/YFP)Amc*^) show cartilage extracellular matrix remodeling, proteoglycan loss and chondrocyte fate alterations, together with an increased expression of chondrocyte hypertrophic markers, as *Col10a1* and *Mmp13*(14), although the characteristic cartilage OA lesions were not observed. On the contrary, Smo genetic downregulation (*COL2-rtTA-Cre;Smo*^*tm2Amc*^) and conditional deletion (*Rosa-CreER(T);Smo*^*fl/fl*^), and *Ihh* conditional deletion (*Col2a1-CreER*^*T2*^*;Ihh*^*fl/fl*^) attenuate surgically induced OA in mice(14,18,19). The effect of Hh pharmacological inhibition employing SMO and GLI inhibitors, such as cyclopamine (CPA) and GANT-61, has also been tested. The treatment with the Hh inhibitor C_31_H_42_N_4_O_5_prevents joint destruction in OA mice, and was associated with a decrease in OA markers, as *Adamts5* and *Col10a1*(14). The efficacy of these inhibitors on cartilage damage attenuation has also been tested in rodent models of severe OA, or adjuvant-induced arthritis(20,21).

On the other hand, blocking of SMO function results in a robust inhibition of Hh signaling(22,23). Potent SMO and GLI antagonists are particularly valuable for effectively inhibiting Hh signaling in several types of tumors with aberrant Hh activation, such as basal cell carcinoma and medulloblastoma(24). However, the accompanying adverse reactions, such as weight loss, fatigue, muscle spasms, alopecia, and dysgeusia(25), makes it questionable to consider SMO inhibitors as a feasible therapy for a chronic joint disease like OA.

Unlike other proteins of the Hh pathway, Evc works as a modulator of Hh signaling that lacks the critical role of other mediators of the Hh pathway, as SMO. In fact, a proportion of *Evc*^*-/-*^ mice in a C57BL/6J;129 mixed background were found to be able to survive for at least 18 days after birth, albeit exhibiting severe skeletal defects(26), whereas *Smo*^*-/-*^ mice do not survive beyond 9.5 days of embryonic development(27). This suggests that Evc silencing may result in a partial blockade of Hh signaling, and may represent a more plausible therapeutic target for Hh inhibition and downstream gene repression in OA cartilage.

Thus, we hypothesized that *Evc* deletion would prevent chondrocyte hypertrophy associated to OA. In order to test whether Evc should be considered as a new therapeutic target in OA, we used our previously reported *Evc* tamoxifen (TAM) induced conditional knockout (*Evc*^*cKO*^) model(28) to specifically study if the blockade of Hh signaling and the entailing hypertrophy mechanisms exclusively during the adult stage could prevent the development of OA.

## RESULTS

### *Evc* levels are drastically diminished in *Evc*^*flox/-*^ mice

Prior the study of *Evc* deletion on OA cartilage damage *in vivo*, we wanted to verify the efficacy of TAM on the deletion of *Evc* in adult *Evc*^*cKO*^ mice. We used RT-qPCR to study *Evc* transcript levels in mice treated with TAM (*Evc*^*cKO*^) and with vehicle (*Evc*^*flox/-*^) as well as in WT mice, which were used as the reference group for normal *Evc* gene expression levels in healthy status. As predicted, *Evc*^*cKO*^ mice did not express *Evc* in lung, heart, brain, muscle or bone tissue (Sup.Fig.1). Unexpectedly, *Evc* levels in *Evc*^*flox/-*^mice were drastically decreased with respect to WT mice, being more similar to those observed in *Evc*^*cKO*^ animals (Sup.Fig.1). For this reason, WT mice, instead of *Evc*^*flox/-*^, were selected as the control group to study the effect of *Evc* deletion on OA cartilage damage *in vivo*.

### *Evc* deletion in DMM-*Evc*^*cKO*^ mice does not prevent OA-associated cartilage damage

We first studied cartilage damage in mice following 8 weeks post-surgery. Experimental OA due to knee joint destabilization evoked articular cartilage lesions in DMM-WT and DMM-*Evc*^*cKO*^ mice compared with their respective healthy controls(28). However, we found no amelioration in cartilage damage of DMM-*Evc*^*cKO*^ mice compared with DMM-WT animals, as showed by similar OARSI scores in both groups (Table 1).

**Table 1.** Histopathological cartilage score in mouse knee joints. OARSI score in NO-WT, DMM-WT and DMM-*Evc*^*cKO*^ mouse knee joints. Data are expressed as mean ± SEM (NO-WT n=10; DMM-WT n=11; DMM-*Evc*^*cKO*^ n=11).*p<0.05 vs NO-WT, ^#^p<0.05 vs DMM-WT.

| **Table 1** Histopathological score in mouse knee joints |  |  |  |
| --- | --- | --- | --- |
| Group | NO-WT | DMM-WT | DMM-*Evcc*<sup>KO</sup> |
| OARSI Score (SEM) | 1.15 (0.198) | 2.818 (0.615)* | 2.727 (0.718) |

### Hh signaling is effectively blocked in DMM-*Evc*^*cKO*^ mice

Meniscal destabilization-induced OA was responsible for an induction in Hh related genes – *Ptch1, Gli1, Evc* and *Ihh* – in the knees of DMM-WT mice in comparison with NO-WT healthy individuals, while the expression of these genes was decreased in DMM-*Evc*^*cKO*^ mice compared with DMM-WT (Fig.1A-D). These results indicate that *Evc* deletion prevents Hh overexpression associated to OA.

**Figure 1.**
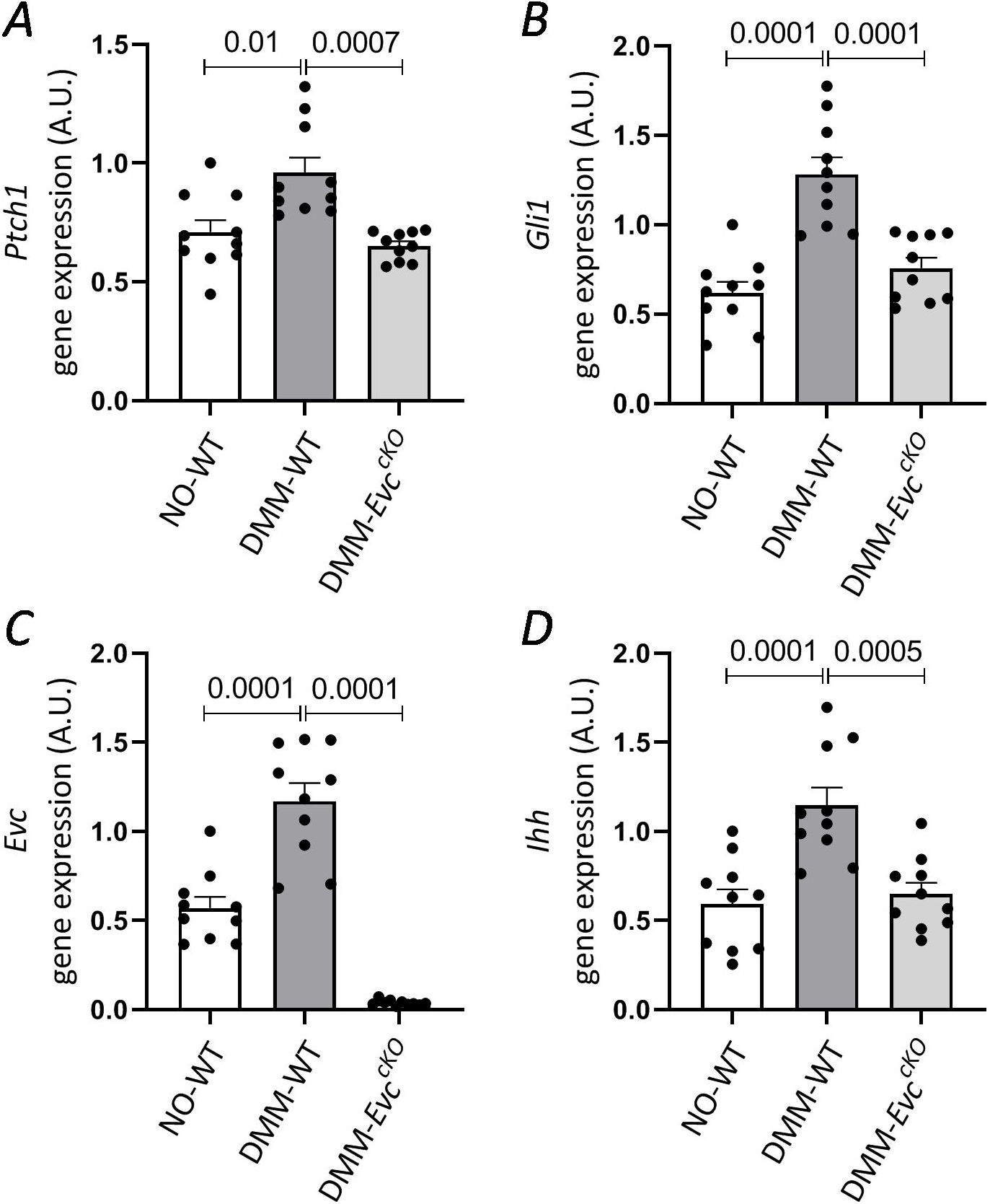
Gene expression of Hedgehog (Hh) mediators in the OA *Evc*^*cKO*^ model and cartilage structure. Gene expression of *Ptch1* (A), *Gli1* (B), *Evc* (C) and *Ihh* (D) in the knees of NO-WT, DMM-WT and DMM-*Evc*^*cKO*^ mice. Data are represented as the individual measurements of each joint and expressed as mean ± SEM (NO-WT n≥7; DMM-WT n≥7; DMM-*Evc*^*cKO*^ n≥7).

### *Evc* deletion does not prevent cartilage catabolism in DMM-*Evc*^*cKO*^ mice

We studied MMP protein profile in the mouse knees to further assess tissue damage in the joint and to determine if *Evc* deletion could attenuate cartilage catabolism in DMM-*Evc*^*cKO*^ mice. MMP-13, MMP-1 and MMP-3 protein levels increased in the knee joints of DMM-WT mice compared with NO-WT, while DMM-*Evc*^*cKO*^ mice showed similar MMP levels compared with DMM-WT animals (Fig.2A-F). This suggests that despite Hh blockade mediated by *Evc* inactivation, cartilage catabolism remains strongly active in DMM-*Evc*^*cKO*^ mice. The analysis of anabolic markers revealed substantial low levels of *Col2a1* gene expression in DMM-*Evc*^*cKO*^ mice, whereas *Agg* transcript levels were similar between groups (Fig.2G,H).

**Figure 2.**
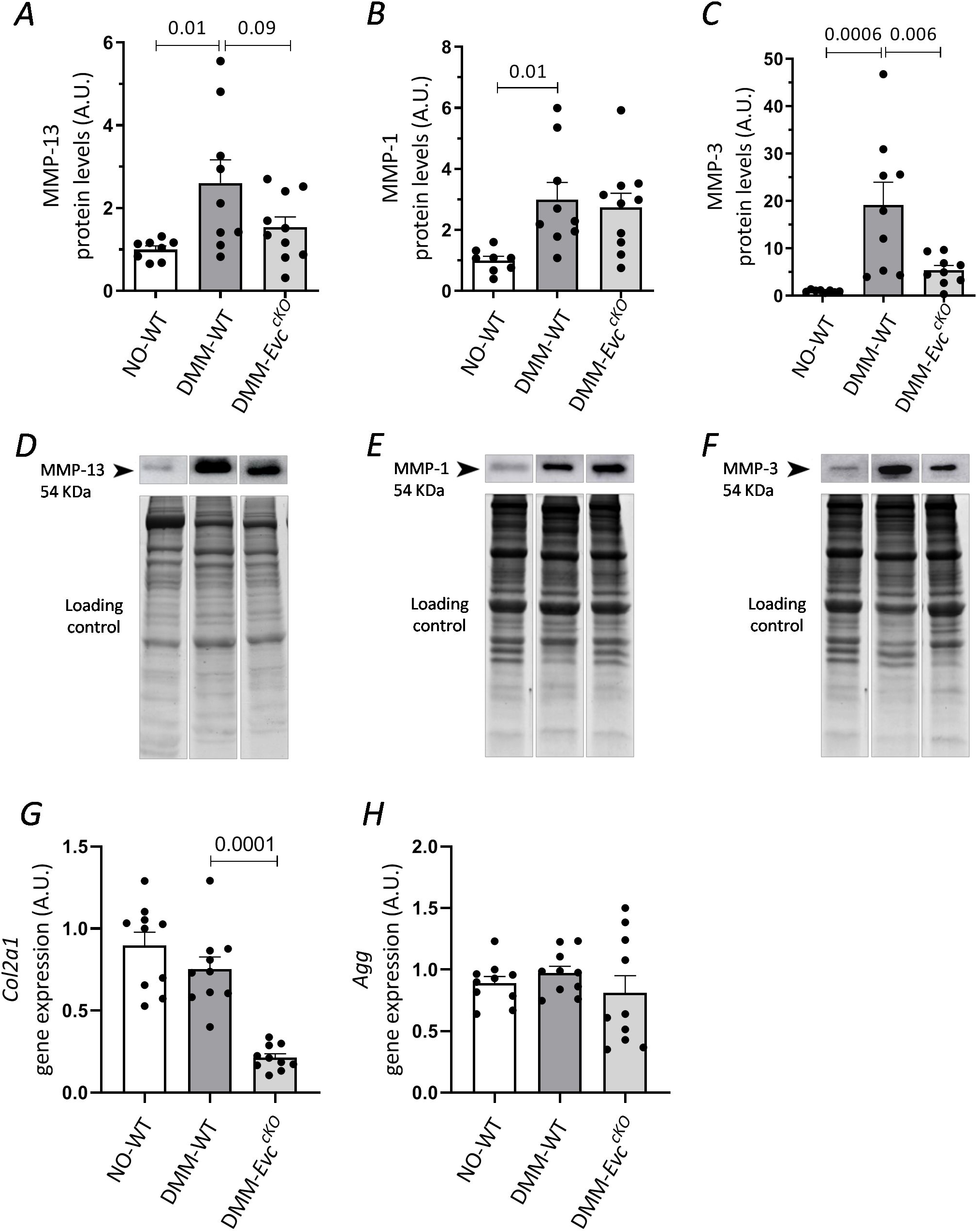
Metalloproteinases (MMP) protein levels in mouse knee joints. Protein levels of MMP-13 (A), MMP-1 (B) and MMP-3 (C) in the knees of NO-WT, DMM-WT and DMM-*Evc*^*cKO*^ mice and their representative western blots (D,E,F). Gene expression of *Col2a1* (G) and *Agg* (H), in the knees of NO-WT, DMM-WT and DMM-*Evc*^*cKO*^ mice. Data are represented as the individual measurements of each joint and expressed as mean ± SEM (NO-WT n≥7; DMM-WT n≥7; DMM-*Evc*^*cKO*^ n≥7).

### Chondrocyte hypertrophy is partially inhibited in DMM-*Evc*^*cKO*^ mice

The analysis of the gene expression levels of hypertrophic markers in the knee joints revealed increased levels of *Ihh, Col10a1* and *Alpl* during OA in DMM-WT mice, which decreased in the DMM-*Evc*^*cKO*^ group with respect to DMM-WT animals (Fig.1D, Fig.3B,E). *Runx2* and *Sp7* levels were also diminished in DMM-*Evc*^*cKO*^ mice compared with DMM-WT (Fig.3A,F). *Mmp13* and *Adamts5* mRNA levels were not modified between groups (Fig.3C,D).The thickness of the femur calcified cartilage showed an increasing trend in DMM-WT animals with respect to NO-WT mice, and decreased in DMM-*Evc*^*cKO*^ mice compared with the DMM-WT group (Fig.3G,H). These results suggest a relevant role of Evc in the hypertrophy and calcification process associated to OA in this model.

**Figure 3.**
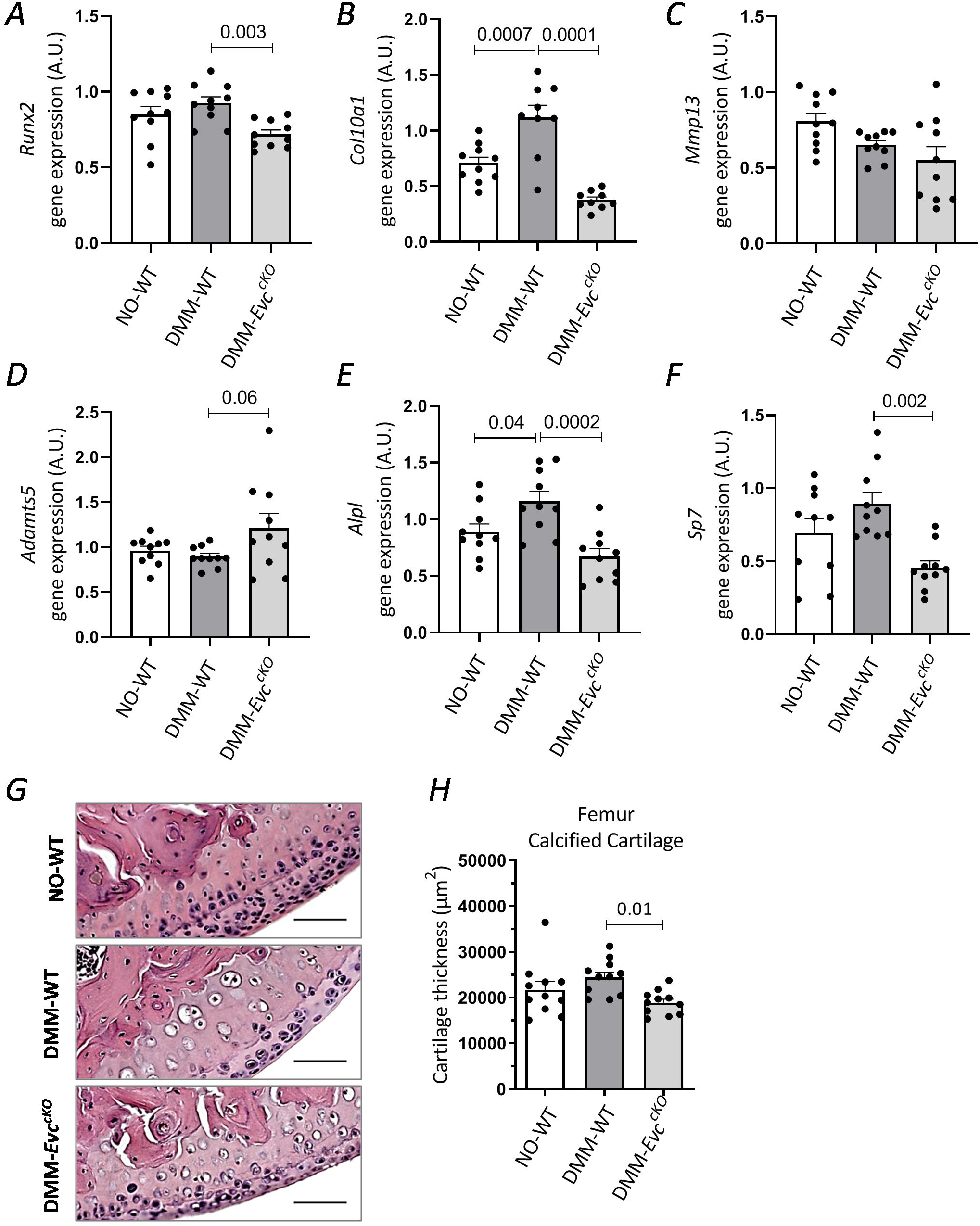
Effect of *Evc* deletion on OA-associated chondrocyte hypertrophy *in vivo*. Gene expression of chondrocyte hypertrophic markers *Runx2* (A), *Col10a1* (B), *Mmp13* (C) and *Adamts5* (D), *Alpl* (E) and *Sp7* (F) in the knees of NO-WT, DMM-WT and DMM-*Evc*^*cKO*^ mice. Representative femur cartilage sections stained with Hematoxylin/Eosin (G). Cartilage thickness in femur calcified cartilage of NO-WT, DMM-WT and DMM-*Evc*^*cKO*^ mice (H). Data are represented as the individual measurements of each joint and expressed as mean ± SEM (NO-WT n≥7; DMM-WT n≥7; DMM-*Evc*^*cKO*^ n≥7).

### Human OA cartilage co-expresses hypertrophic and inflammatory phenotypes

We suspected that the blockade of the hypertrophy response could trigger a higher inflammatory activation in OA chondrocytes. This would explain the fact that despite blocking hypertrophy we did not see improvement in cartilage damage in DMM-*Evc*^*cKO*^ mice. To test this premise, we first aimed to determine whether chondrocytes polarize towards the acquisition of a hypertrophic vs an inflammatory phenotype in OA cartilage. With this purpose we analyzed the presence of IHH and COX-2 in human cartilage samples by immunofluorescence (Fig.4A,B). We found that both proteins co-localized in the same cells and areas in human OA cartilage (Fig.4D), suggesting that chondrocytes can express both hypertrophic and inflammatory proteins at the same time. This result demonstrates the coexistence of both pathological phenotypes in the same cell.

**Figure 4.**
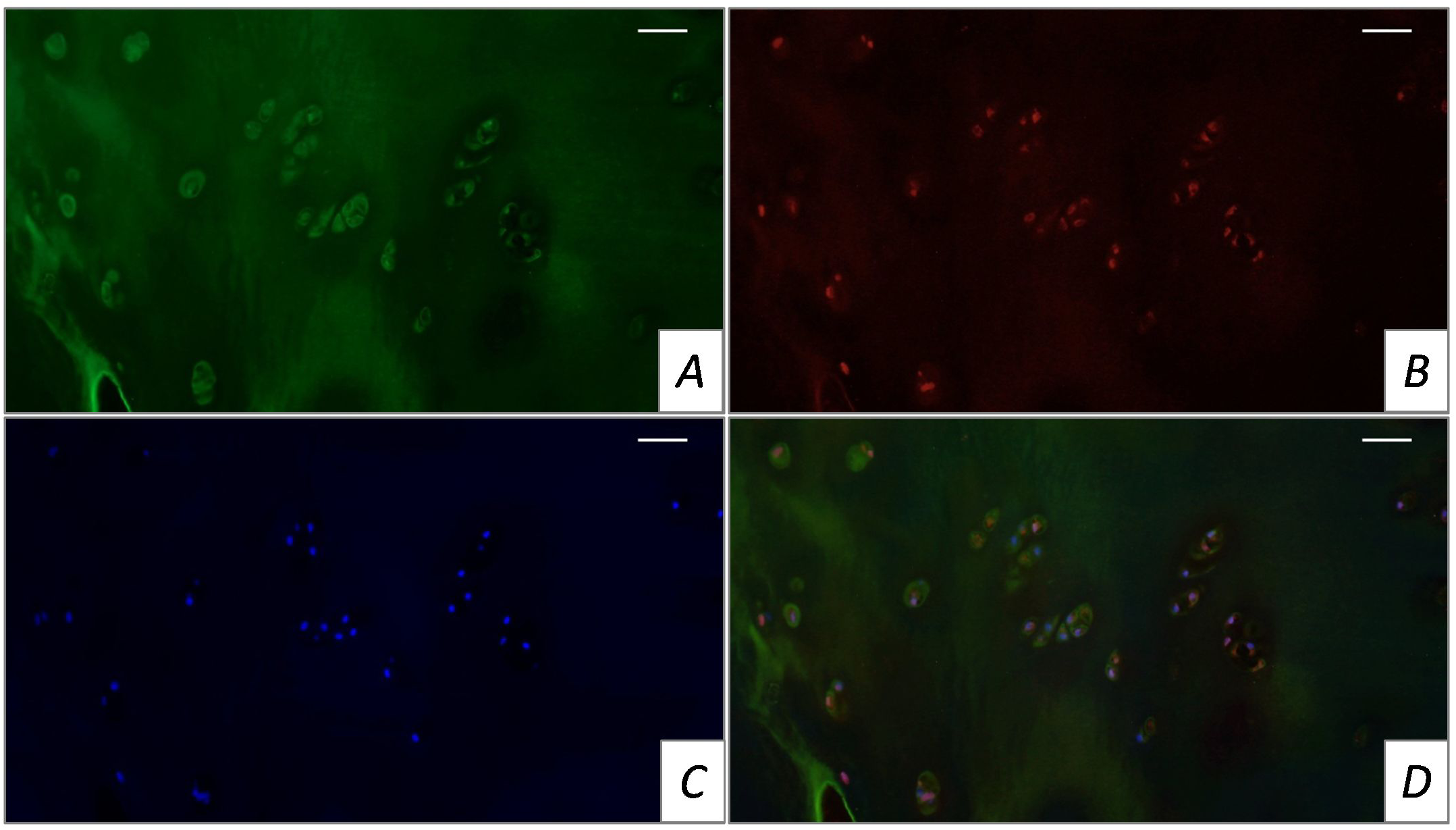
Co-localization of hypertrophic and inflammatory markers in human cartilage. Immunofluorescence of IHH (A) and cyclooxygenase-2 (COX-2) (B), chondrocyte nuclei staining with DAPI (C)and merge (D) in human OA cartilage samples.

### Human OA chondrocytes inflammatory response is not modified by Hh inhibition

Once we observed that both hypertrophic and inflammatory phenotypes co-localize in the same cell, we aimed to determine whether both signaling processes interact and mutually regulate in OA chondrocytes. We studied if IL-1 beta-mediated inflammatory response could indirectly modify the synthesis of hypertrophic differentiation inducers in human OA chondrocytes. IL-1 beta insult decreased *GLI1* and *EVC* gene expression levels in IL-1 beta-stimulated chondrocytes compared with control cells (Fig.5B,C), and increased *IHH* gene expression levels (Fig.5D), while no changes were observed in *PTCH1* expression (Fig.5A). Then, we deepened in the inflammatory study in human OA chondrocytes to determine the effect of Hh signaling blockade on the response of chondrocytes to IL-1 beta-induced inflammation. We used CPA as a Hh inhibitor due to its direct antagonism by interaction with SMO(29). CPA-mediated Hh inhibition did not modify the gene expression of pro-inflammatory markers – *IL1B, MMP13, PTGS2, PTGER2, IL6, NOS2* and *CCL2* – with respect to chondrocytes not treated with the SMO inhibitor (Fig.5E-K). These data indicate that Hh inhibition, and therefore hypertrophy blockade, does not alter OA chondrocytes inflammatory response to IL-1 beta.

**Figure 5.**
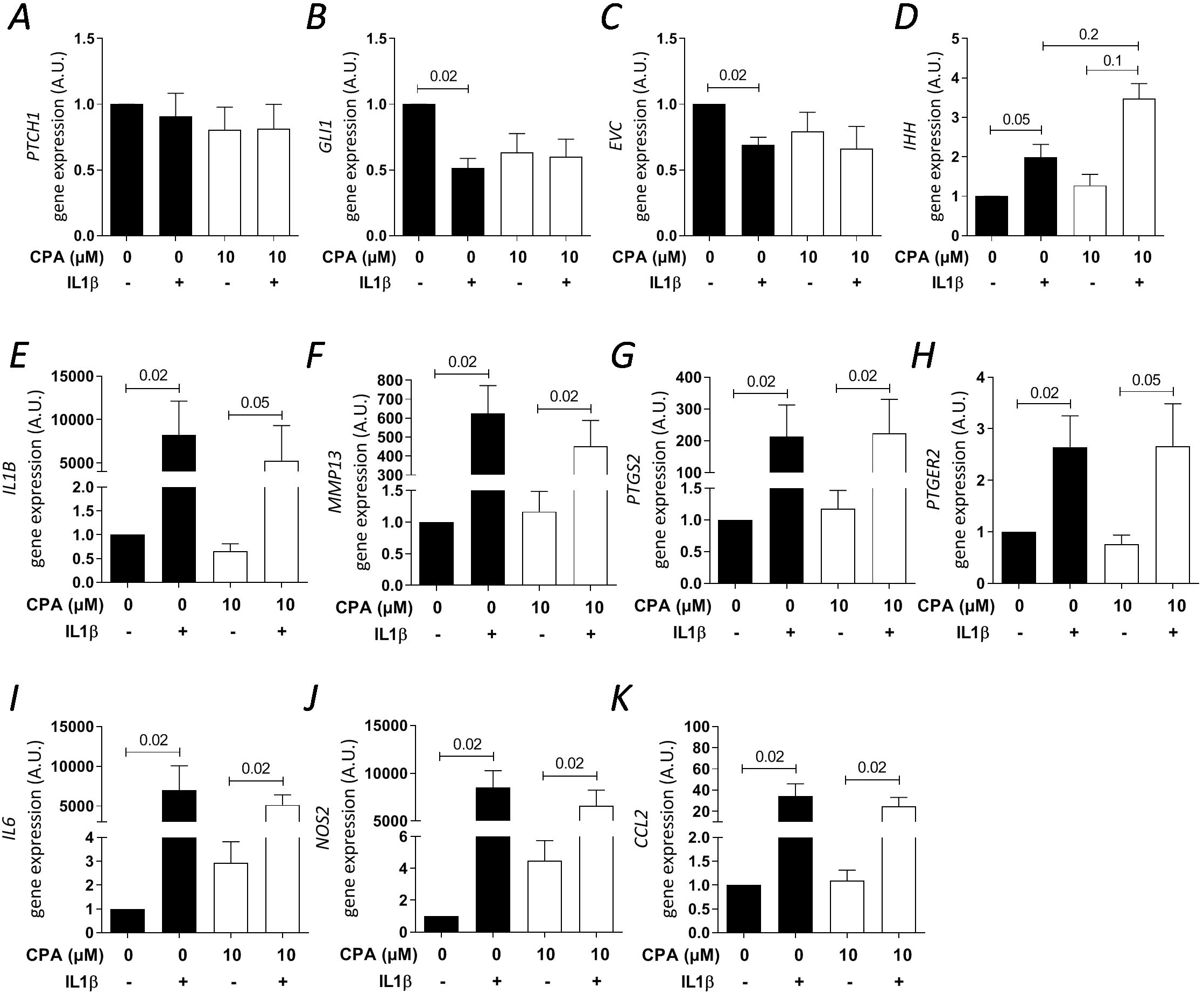
Inflammatory effect of IL-1 beta on human OA chondrocytes *in vitro*. Gene expression of Hh signaling mediators *PTCH1* (A), *EVC* (B), *GLI1* (C) and *IHH* (D) and proinflammatory mediators *IL1B* (E), *MMP13* (F), prostaglandin G/H synthase 2 (*PTGS2)* (G), prostaglandin E2 receptor (*PTGER2*) (H), *IL6* (I), inducible nitric oxide synthase (*NOS2*) (J) and monocyte chemoattractant protein-1 (*CCL2*) (K) in human OA chondrocytes treated with IL-1 beta and CPA, or vehicle (DMSO), for 24 hours. Data are normalized by chondrocytes in basal conditions (no IL-1 beta, no CPA stimulation) and expressed as mean ± SEM (n=5 independent experiments).

## DISCUSSION

Our results show that surgically-induced OA in mice induced the activation of the Hh pathway, increasing *Ptch1, Gli1* and *Ihh* gene expression in the OA knees, as previous studies have observed in rodent models of OA(14,30). In addition, we observed that *Evc* levels were also increased in OA mouse knee joints.

The role of EVC and its specific activation have not been previously described in OA. Since EVC positively mediates Hh signaling, and DMM induces its overexpression, we proposed EVC as a new therapeutic target for inhibiting chondrocyte hypertrophy-like phenomena that are activated in OA. In addition, the expression of *Evc* seems to be mainly limited to cartilage tissue(26,31). Thus, a selective EVC inhibition would be specifically directed to articular hyaline cartilage in comparison to different Hh inhibitors that have a widespread effect in different tissues and organs.

Hh signaling has an essential role in limb and neural development and is dysregulated in disease states, such as in genetic (e.g. Bardet–Biedl syndrome) or neurodegenerative disorders (e.g. Parkinson’s and Alzheimer’s disease), cancer (e.g. basal cell carcinoma (BCC) and medulloblastoma), and OA(32,33). Hh function is also required for tissue repair and homeostasis, in coordination with WNT and TGF-beta/BMP signaling and G protein-coupled receptors actions(33), for example in lung epithelium regeneration(34), epidermal and hair follicle maintenance(35), and muscle regeneration(9).

The mechanisms of Hh signaling inhibition at different levels of the pathway have been deeply studied in the context of cancer research. Different Hh pharmacological inhibitors are available for cancer treatment, mainly SMO and GLI1 inhibitors, which effectively suppress Hh activation(36,37). These Hh-targeted drugs have shown their efficacy in the treatment of a variety of tumors, including BCC, pancreatic or breast cancer(37). However, these Hh inhibitors, like vismodegib and sonidegib, are associated with different adverse events such as muscle spasms, alopecia, dysgeusia, weight loss, and fatigue(25,38).

The experimental use of these inhibitors in OA models has proven that SMO and GLI pharmacological inhibitors strongly restrain Hh signaling and exert a protective effect against OA progression(14,39,40). OA mice treated with the SMO inhibitor C_31_H_42_N_4_O_5_ showed articular cartilage recovery, as well as a downregulation of *Ptch1, Gli1*, and Hh interacting protein (*Hhip*), together with a decrease in the hypertrophic markers *Adamts5* and *Col10a1*(14). In rats with adjuvant-induced arthritis, CPA reduced cartilage damage and inflammation, diminishing TNF-a, IL-1 beta, and IL-6 serum levels(21). Furthermore, the GLI inhibitor GANT-61, in combination with a low dose of the anti-inflammatory indomethacin, was found to synergistically reduce cartilage damage and inflammatory cytokines TNF-a, IL-2, and IL-6 in serum, through pyroptosis inhibition in chondrocytes(20).

Although these Hh inhibitory molecules have been tested in experimental OA models and have given promising results, the use of these compounds for the treatment of human OA does not seem acceptable due to their toxicity and the above mentioned associated adverse effects. Also, their long-term effects on the arthritic joint itself, and in other regenerative processes such as bone fracture healing, are still undefined. Due to the less critical nature of Evc in comparison to other mediators of the Hh pathway, Evc inhibition would provide a more plausible target for the treatment of OA. Although half of *Evc* knockouts were reported to die soon after birth, some can survive to adulthood under special heed in a C57BL/6J;129 mixed genetic background. In contrast, *Smo* knockouts fail embryonic development and die at E9.5(26,32,41). Thus, Evc blockade would display a partial inhibition of the Hh pathway in comparison with the vast inhibition induced by the blockade of other mediators, such as SMO or IHH.

Aside from the classical Hh activation, several pathways such as MAPK/MEK/ERK, PI3K/AKT/mTOR, TGF-beta and PKC signaling, modulate GLI1 and GLI2 transcriptional activity through non-canonical Hh activation(42). Particularly in OA, Hh signaling interacts with the NOTCH, WNT, FGF and mTOR pathways(2). Furthermore, inflammatory pathways such as TNF-a/mTOR, potentiate GLI1 activity in a SMO-independent manner(43,44).Overall, signaling cascades and cellular responses triggered by Hh pathway seem to be highly context-dependent(2).

We utilized an *Evc*^*cKO*^ model in mice females combined with DMM, and we demonstrated that *Evc*^*cKO*^ effectively inhibited Hh signaling overexpression during OA triggered by DMM, and decreased chondrocyte hypertrophic markers. We employed this OA model, in which mice developed mild to moderate OA alterations(45), in order to test plausible beneficial effects of hypertrophy blockade. We observed a general decrease in the expression of *Ihh, Runx2, Col10a1, Alpl* and *Sp7* in the joints of DMM-*Evc*^*cKO*^ mice in comparison to DMM-WT animals. Furthermore, tamoxifen induced inactivation of Evc accounted for a substantial prevention of calcified cartilage thickening in DMM-*Evc*^*cKO*^ mice compared to DMM-WT mice.

Although *Evc*^*cKO*^ showed a decrease in Hh induction induced by OA, together with a lower hypertrophy and calcification progression, no amelioration of cartilage damage was observed in DMM-*Evc*^*cKO*^ mice compared with DMM-WT. We have previously demonstrated that DMM-WT and DMM-*Evc*^*cKO*^ mouse joints also exhibited similar subchondral bone sclerosis associated to DMM(28).

Research in *Evc* null mice have shed light on its physiological role and demonstrated that EVC localises at the base of the primary cilium and mediates Hh signaling in chondrocytes and osteoblasts(26,46). Still, the role of Evc in the cartilage seems to be restricted to skeletal development and bone growth, with an unclear role in the adult tissue. Specifically in OA, studies have been conducted investigating the function of the primary cilium in chondrocytes. An increased cilium lengthand prevalence of ciliated chondrocytes have been associated to tissue erosion and mild to severe OA lesions in the cartilage(47,48). Also cilia orientation undergoes alterations in the OA cartilage. While healthy chondrocytes orient their cilium away from the cell surface, chondrocytes in OA cartilage direct their primary cilia into the core of OA chondrons(48). Alterations in primary cilium structure and assembly have also been associated with a Hh-mediated mechanosensitive diminished response in bovine articular chondrocytes(49). Similarly, primary cilium depletion in chondrocytes is responsible for impeded GLI3 processing to the repressor form of the transcription factor, thus promoting increased Hh signaling and OA markers(50). *Evc*^*-/-*^ chondrocytes do not show alterations in ciliogenesis(26). Yet, *Evc* absence does produce chondrocyte dysfunction, as can be asserted by the severe skeletal and growth alterations of *Evc*^*-/-*^and growth plate column disarrangement in *Evc*^*cKO*^ adult mice(26,28). It still remains unknown how inactivation of Evc in *Evc*^*cKO*^ mice might influence articular cartilage quality in the long term.

The extremely low *Col2a1* transcript levels found in DMM-*Evc*^*cKO*^ mice are consistent with the alteration of the genetic locus of *COL2A1* in patients with osteochondrodysplasias(51). In an Ellis–van Creveld experimental model in calf with an *EVC2* mutation, abnormal COL2A1 expression was histologically found in the physis, attributed to an accelerated COL2A1 degradation(52). In contrast, our data show a COL2A1 synthesis defect in DMM-*Evc*^*cKO*^ mice. The low *Col2a1* transcript levels, together with the relatively high MMP-associated catabolism, particularly MMP-13 and MMP-1, maintained in the knee joints of DMM-*Evc*^*cKO*^ mice, may have contributed to a more rapid progression of cartilage damage in DMM-*Evc*^*cKO*^ mice than a priori anticipated. Both findings matched histological cartilage damage in DMM-*Evc*^*cKO*^ joints and its equal OA progression to DMM-WT mice(28).

Both hypertrophy-like alterations and inflammation in OA chondrocytes have been described as characteristic phenomena in this disease. Recent data from our laboratory demonstrate that hypertrophic chondrocytes can be found in all hyaline cartilage layers, and not within a specific location(16). In this work, we have demonstrated that these signals can be simultaneously found in OA chondrocytes, since we were able to co-localize IHH and COX-2 in human OA chondrocytes. The co-localization of both hypertrophy and inflammatory markers in human OA cartilage suggests that the acquisition of the hypertrophic-like phenotype does not exclude the expression of an inflammatory profile. Our data indicate the presence of a complex signaling network modulating chondrocyte responses in OA.

In line with this data, it could be possible that IHH-signaling down-regulation observed in *Evc*^*cKO*^ mice could exacerbate the inflammatory response in OA chondrocytes, being responsible for the absence of a protective effect on cartilage degradation observed in DMM-*Evc*^*cKO*^ mice, in comparison to DMM-WT. In fact, it has been previously shown that both pathways could be able to regulate each other. IL-1 beta stimulation would decrease the activation of the Hh pathway(53), while Hh activation has been long associated with OA cartilage degradation due to an increase in catabolic enzymes such as ADAMTS5 and MMP-13(13,14,18). However, Thompson and co-workers observed that Hh pathway activation induced by recombinant-Ihh did not stimulate cartilage degradation in healthy bovine articular chondrocytes(53). On the other hand, some reports have even postulated that Hh may induce anti-inflammatory responses(54,55). Genetic or pharmacologic Hh inhibition increased the presence of inflammatory mediators, whereas *Gli1* overexpression seemed to ameliorate inflammatory outcomes in mouse models of colitis and acute pancreatitis through the induction of the anti-inflammatory cytokine IL-10(54,55).

Thus, employing *in vitro* studies, we investigated whether Hh signaling inhibition might trigger a higher inflammatory response in human OA chondrocytes. Our data indicated that IL-1 beta accounted for an inhibition in Hh pathway in human OA chondrocytes, particularly decreasing *GLI1* and *EVC* gene expression. In contrast, we observed that human OA chondrocytes did not modify their inflammatory response to IL-1 beta under Hh pathway inhibition.

Overall, our results showed that *Evc*-mediated Hh inactivation partially prevented chondrocyte hypertrophy but did not ameliorate OA cartilage damage in DMM-*Evc*^*cKO*^ mice. These data suggest that a partial blockade of the Hh pathway through tamoxifen induced inactivation of *Evc* is not a therapeutic target for mild/incipient OA. In this OA model, we have demonstrated that chondrocyte hypertrophy was associated to Hh signaling activation, but it is not a pathogenic event in the development of the disease. In this sense, chondrocyte hypertrophy could be a frustrated regenerative mechanism that correlates with OA progression, but not a leading cause of cartilage degeneration per se.

## MATERIALS AND METHODS

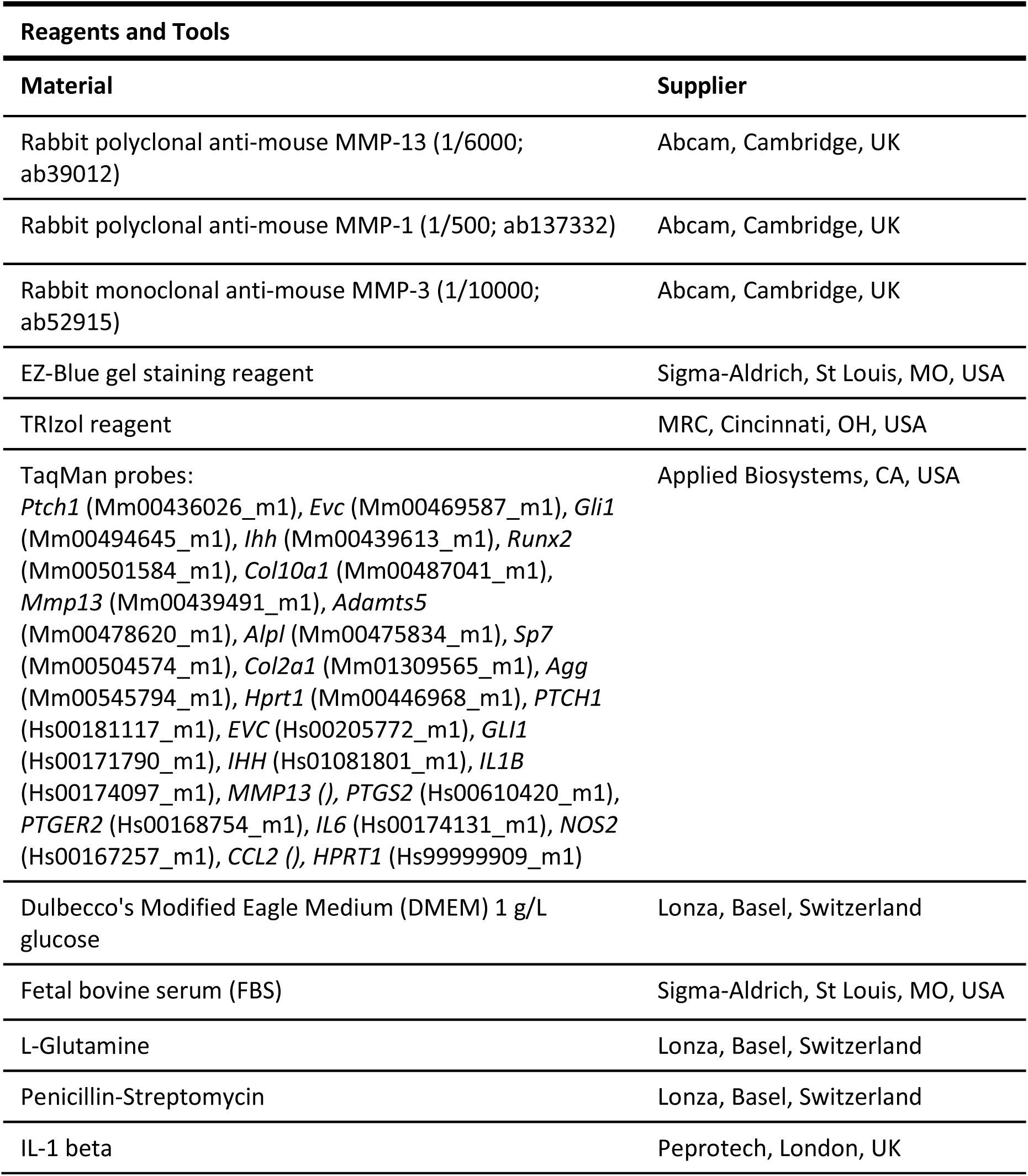
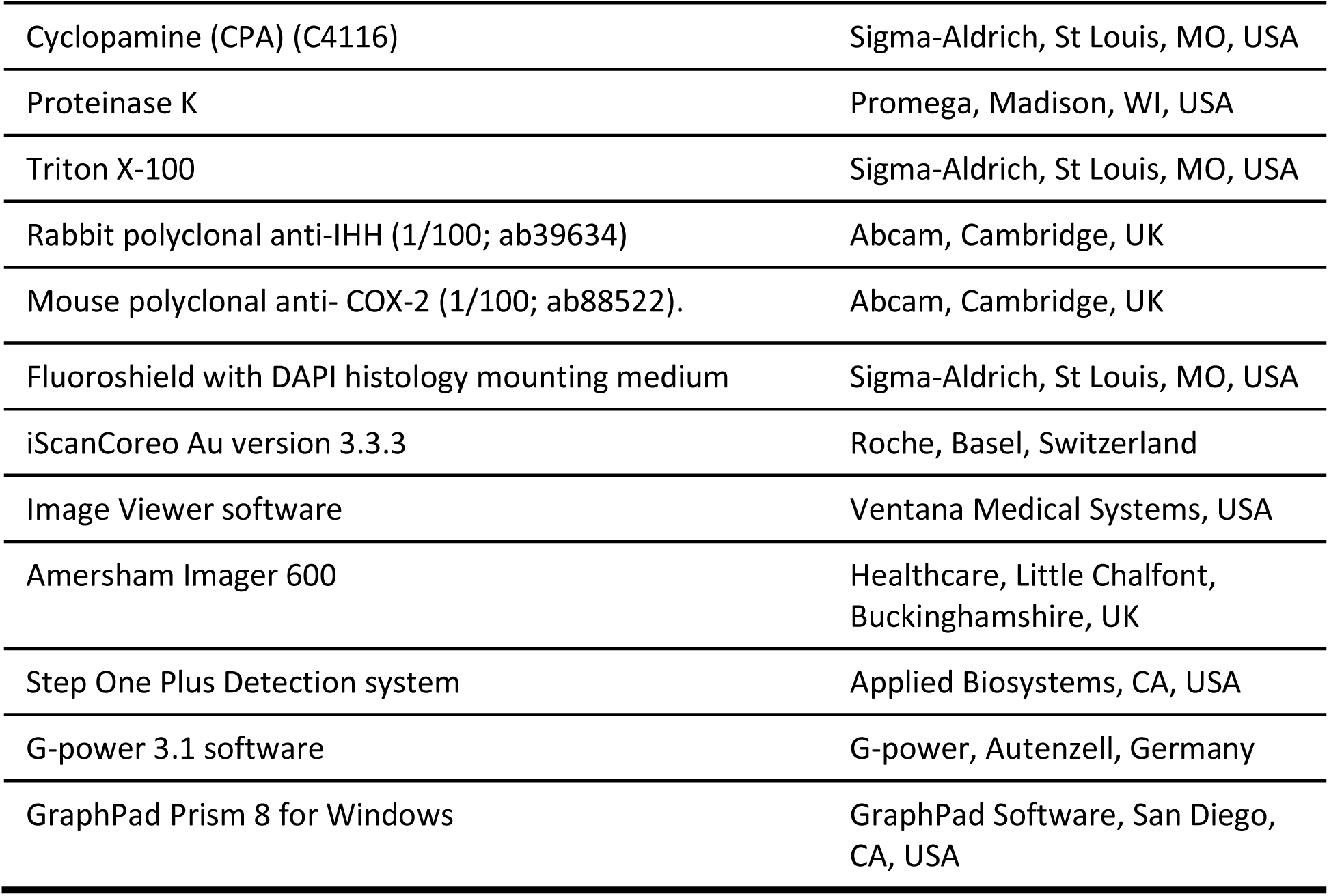

### Methods and Protocols

#### Animal model of OA

Generation of *Evc*^*flox/-*^; UBC-*CreERT2* mice was previously described(28). These mice were maintained in a C57BL/6J;129 mixed background. Briefly, 10 weeks-old *Evc*^*flox/-*^;*UBC-CreERT2*^*+/-*^ female mice received vehicle or 0.075 mg/g/day TAM administered in five intraperitoneal injections. Vehicle-treated mice were named as *Evc*^*flox/-*^ and TAM-treated mice as *Evc*^*cKO*^. C57BL/6J;129 WT mice were used as controls due to unexpected low *Evc* levels in *Evc*^*flox/-*^mice (Sup.Fig.1). Mice were separated and housed in cages according to their genotype to avoid tamoxifen contamination, were exposed to 12-hour light/dark cycles, and had free access to water and standard chow. OA was induced at 12 weeks of age by destabilization of the medial meniscus (DMM) as previously described(28,45), and mice were assigned to groups of study according to their genotype and randomly distributed between healthy and DMM-operated: non-operated (NO)-WT (n=8), DMM-WT (n=6)and DMM-*Evc*^*cKO*^ (n=6). After 8 weeks of OA progression, mice were euthanized and tissues and joints collected for histopathological and molecular analysis. Tissue samples from brain, lung, muscle, heart, bone and joints were immediately frozen for molecular biology studies. Animal handling and experimental procedures for this study (PROEX 119/16) complied with the national and international regulations, and the Guidelines for the Care and Use of Laboratory Animals (NIH), and were approved by the Institutional Ethics and Welfare Committees of the IIS-Fundación Jiménez Díaz and Alberto Sols Biomedical Research Institute.

#### Cartilage thickness analyses

Cartilage thickness was evaluated in knee sections stained with Hematoxylin/Eosin and scanned with the iScanCoreo Au version 3.3.3 (Roche, Basel, Switzerland). A 500 μm line was drawn covering the center and posterior region of the joint in the femur using the Image Viewer software (Ventana Medical Systems, USA), and calcified cartilage area was calculated using the Image J software.

#### Western blotting

20 μg of protein extract from knee joints were separated by SDS-PAGE and transferred to nitrocellulose membranes as previously described(56,57). Membranes were incubated overnight at 4°C with the following antibodies: rabbit polyclonal anti-mouse MMP-13 (1/6000; ab39012; Abcam) and MMP-1 (1/500; ab137332; Abcam), and rabbit monoclonal anti-mouse MMP-3 (1/10000; ab52915; Abcam). Binding signal was detected with chemiluminescence in an Amersham Imager 600 (Healthcare, Little Chalfont, Buckinghamshire, UK). EZ-Blue gel staining reagent (Sigma) was used as loading control and densitometric measures were normalized by the average value of the NO-WT and expressed as arbitrary units (A.U.)(56–58).

#### RNA isolation and gene expression assays

RNA was isolated from frozen knees, prior crush in liquid nitrogen, by TRIzol reagent (MRC, Cincinnati, OH, USA) and retrotranscribed to cDNA as described elsewhere(56). RNA from chondrocyte cultures was also isolated with TRIzol reagent. Gene expression was quantified by real-time PCR using the Step One Plus Detection system (Applied Biosystems, CA, USA). TaqMan probes for mouse *Ptch1, Evc, Gli1, Ihh, Runx2, Col10a1, Mmp13*, A disintegrin and metalloproteinase with thrombospondin motifs 5 (*Adamts5)*, Alkaline phosphatase (*Alpl)*, Transcription factor SP7 (*Sp7)*, type II collagen (*Col2a1*) and aggrecan (*Agg*), and TaqMan probes for human *PTCH1, EVC, GLI1, IHH, IL1B, MMP13*, prostaglandin G/H synthase 2 (*PTGS2*), prostaglandin E2 receptor (*PTGER2*), *IL6*, inducible nitric oxide synthase (*NOS2*) and monocyte chemoattractant protein-1 (*CCL2*) were purchased from Applied Biosystems. Gene expression levels were determined with the comparative Ct quantitation method using Hypoxanthine-guanine phosphoribosyltransferase (*Hprt1, HPRT1)* as internal control and expressed as arbitrary units (A.U.).

#### Human OA cartilage collection for chondrocyte isolation

Human OA cartilage was obtained from patients undergoing knee joint replacement surgery (Fundación Jiménez Díaz Hospital), prior informed consent and approval from the Institutional Ethics Committee, and following the ethical principles of the Declaration of Helsinki and the Department of Health and Human Services Belmont Report. Chondrocytes were isolated as previously described(59).

#### Culture of human chondrocytes

Human OA chondrocytes were seeded at a confluence of 2·10^5^ cells/well in p6 plates with Dulbecco’s Modified Eagle Medium (DMEM) 1 g/L glucose, supplemented with 10% fetal bovine serum (FBS)(Sigma), 2 mM glutamine (Lonza), and 100 U/ml Penicillin-Streptomycin (PS) (Lonza). Experiments were performed in passage 2. Cells were FBS-depleted for 24 hours, and then stimulated with 1 ng/mL IL-1 beta (Peprotech, London, UK), and 10 μM CPA (Sigma), a direct Hh antagonist(29), for 24 hours. Chondrocytes not stimulated with IL-1 beta nor CPA were used as control. Each experiment was performed with chondrocytes from different donors.

#### Immunofluorescence of human cartilage

Immunofluorescence was performed based on previously described protocols(60). Briefly, 3 μm knee joint sections were deparafinized and rehydrated. Antigen retrieval was performed by incubation with 20 μg/mL proteinase K (Promega, USA) for 20 minutes. Tissue sections were incubated with 0.1 M glycine for autofluorescence removal, and blocked with 3% PBS-bovine serum albumin (BSA), 0.1% Triton X-100 (Sigma), 5% FBS. Then cartilage sections were incubated with the corresponding primary antibodies: rabbit polyclonal anti-IHH (1/100; ab39634, Abcam) and mouse polyclonal anti-Cyclooxygenase-2 (COX-2) (1/100; ab88522; Abcam). Secondary FITC and TRITC respectively antibodies were used for detection of positive fluorescence signal. Tissue sections were ultimately incubated with 0.1% Sudan Black in 70% ethanol and mounted with Fluoroshield with DAPI histology mounting medium (Sigma). Sections were photographed with a MiCom fluorescence microscope equipped with ACT-1 software at ×40 magnification.

#### Statistical analysis

An animal model previously studied(28)was used for the attainment of the present research. Each limb was analyzed as an independent sample. Due to the lack of previous studies of OA in mice with *Evc* deletion, previously published data of an OA model with *Ihh* deletion was used for the calculation of the sample size(18). The sample size was determined with the objective of detecting differences with respect to the articular cartilage damage score between OA and OA *Col2a1-CreERT2; Ihh*^*fl/fl*^ mice. By accepting a significance level (alpha) of 5% assuming a Bonferroni correction, and a statistical power of 80%, a pairwise t-test between conditions required 6 limbs per group to demonstrate an 86% (7 vs 1 damage score with the *OARSI Osteoarthritis Cartilage Histopathology Assessment System* (OOCHAS) scale assuming a standard deviation of 3 and 2 respectively. Sample size estimation was performed using the G-power 3.1 software (G-power, Autenzell, Germany)(61), which indicated no requirement of additional animals to the study in order to detect the expected differences in cartilage damage between groups.

Ordinary one-way ANOVA with Bonferroni post-hoc test was used for comparisons between groups with normal distribution of the data, based on Shapiro-Wilk normality test. Kruskal-Wallis test was used for comparisons between multiple groups where data lacked normality, followed by Dunn’s post-hoc test. A P-value of less than 0.05 was considered statistically significant. Statistical analyses of data were performed using GraphPad Prism 8 for Windows (GraphPad Software, San Diego, CA, USA). Data were expressed as mean with standard error (SEM).

## Acknowledgements

This work was financially supported by grants from the Instituto de Salud Carlos III [PI15/00340, PI16/00065, PI18/00261] and Fondo Europeo de Desarrollo Regional (FEDER) and by grants SAF-2013-43365-R and SAF2016-75434-R to VLR-P. AL and PG were funded by Fundación Conchita Rábago. We would like to thank Dr. David Santamaria for providing UBC-*CreERT2* mice and María Gracia González Bueno for helping to organize mouse crosses and technical work.

## Author contributions

AL, VLR-P, RL and GH-B were in charge of conceptualization, formal analysis, and interpretation of data. VLR-P, GH-B and RL were responsible for the funding acquisition, provision of resources and project administration. AL, PG, LC, VLR-P, AP-C and RL were in charge of in charge of animal methodology design. LC, VLR-P and AP-C generated the genetically modified *Evc*^*cKO*^ model, and AL, PG, SP-N and RL performed the animal procedures. AL and PG contributed to the acquisition of data, investigation and visualization. AL, PG, LC, VLR-P, AP-C, SP-N, AM, GH-B and RL were involved in the writing process and drafting the article and its revision and editing, and approved the final manuscript to be published. GH-B and RL have full access to overall data and take responsibility for the supervision, validation, integrity and accuracy of the data analysis.

---

**Supplementary Figure 1.**
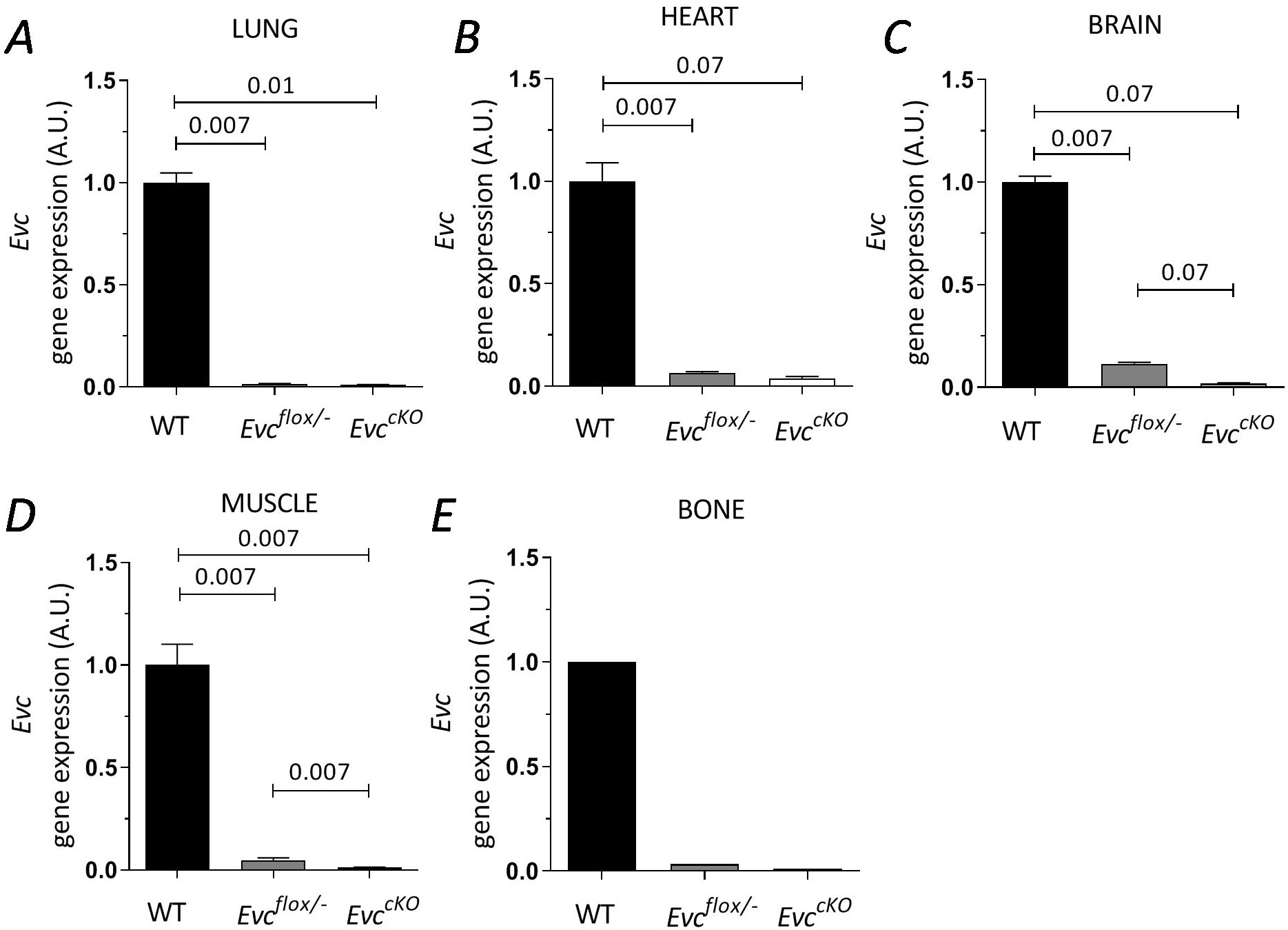
*Evc* mRNA levels in tissues of WT, *Evc*^*flox/-*^ and *Evc*^*cKO*^ animals. *Evc* gene expression in lung (A), heart (B), brain (C), muscle (D) and bone (E) of WT, *Evc*^*flox/-*^ and *Evc*^*cKO*^ mice. Data are normalized with respect to *Evc* mRNA levels in WT mice and expressed as mean ± SEM (WT n=6; *Evc*^*flox/-*^ n = 6; *Evc*^*cKO*^ n≥3 for lung, brain, muscle and heart; WT n=1; *Evc*^*flox/-*^ n = 1; *Evc*^*cKO*^ n = 1 for bone – pool of tibiae –).

